# Shapes of condensate droplets containing filaments

**DOI:** 10.64898/2026.03.31.715246

**Authors:** Fynn Wolf, Shannon Bareesel, Britta J. Eickholt, Roland L. Knorr, Susanna Röblitz, Sushma N. Grellscheid, Halim Kusumaatmaja, Thomas J. Böddeker

## Abstract

The interactions of droplets and filaments can lead to mutual deformations and complex combined behavior. Such interactions also occur within the cell, where biomolecular condensates, distinct liquid phases often composed of proteins, have been observed to structure and affect the organization of the cytoskeleton. In particular, biomolecular condensates have been shown to undergo characteristic deformations when cytoskeletal filaments are fully embedded within them. However, a full understanding of the underlying physical mechanisms is still missing. Here, we combine experiments with coarse-grained molecular dynamics simulations and analytical models to uncover the physical mechanisms that define emerging shapes of droplets containing filaments. We find that the surface tension of the liquid phase and the bending energy of the filament(s) suffice to accurately capture emerging shapes if the length of the filament is small compared to the liquid volume. As the volume fraction of filament(s) increases, wetting effects become increasingly important, setting physical constraints within which surface and bending energies compete to define the droplet shapes. We find that mutual deformations of condensate and filament extend accessible shapes beyond classical stability considerations, leading to structuring and entrapment of contained filaments. Shape deformations may further affect ripening dynamics that favor certain geometries. Our findings provide a physical framework for a better understanding of the possible roles of biomolecular condensates in cytoskeletal organization.

## I. INTRODUCTION

Liquid droplets and filaments are ubiquitous structures in nature. Upon contact, droplet and filament may interact, leading to mutual deformations and complex combined behavior. Such phenomena also occur within the cell, where biomolecular condensates, liquid phases typically composed of proteins and nucleic acids [1], can deform and structure the cytoskeleton through capillary forces [2–4] or affect polymerization kinetics of cytoskeletal filaments by boosting monomer concentrations [5, 6]. In this way, biomolecular condensates may aid or facilitate fundamental biological processes that require (re-)organization of the cytoskeleton, such as structure formation, cellular motility, and mitosis.

We are just starting to uncover the impact of condensates on cytoskeletal organization in the cell. However, interactions between biomolecular condensates and cytoskeletal filaments *in vitro* have been studied in more detail, both for interactions with tubulin/microtubules [7–11] and with actin [5, 12–14], suggesting diverse functions in the cell. Many of these studies report mutual deformations of filament(s) and condensate. A key length scale is the bendocapillary length [15], which balances wetting and bending energy. Small condensates, that are below the bendocapillary length, fail to retain bent filaments, leading to straight filaments wetted by the condensate. For larger condensates, above the bendocapillary length, the adhesion of the filament to the condensate becomes strong enough to bend the filament around the condensate. Moreover, some efforts have been made to formally characterize these deformations using simulations [13, 14, 16, 17] and analytical methods [18, 19]. Nonetheless, a detailed understanding of the interactions between filament and droplets and the resulting different shapes is still missing.

In this paper, we investigate the physical mechanisms that define the shapes of droplets containing entire filaments. We first carry out experiments using biomolecular condensates containing F-actin *in vitro* to characterize the droplet shapes. Based on the experimental results, we develop a simple analytical model that predicts droplet shapes on the assumption that a competition of surface and bending energy is the main contributor. To validate the model and to investigate the droplet-filament system in detail we perform coarse-grained molecular dynamics simulations. These reveal that, particularly in the regime of biomolecular condensates, wetting effects become important, defining boundaries within which surface and bending energies sculpt minimal energy shapes.

## II. RESULTS

### A. Experimental Results

We perform experiments in which we polymerize F-actin inside biomolecular condensates to identify parameters that affect the droplet shapes. The condensate consists of PLPPR3 ICD, which is the intracellular domain of phospholipid phosphatase-related protein 3, a transmembrane protein that regulates filopodia density in developing neurons [20, 21]. PLPPR3 ICD has been shown to form condensates that localize and sustain F-actin polymerization *in vitro* [5]. Condensates are formed at a protein concentration of 20 µM PLPPR3-ICD in buffer that allows for actin polymerization.

Condensates sediment onto a coating of polyvinyl alcohol (PVA) with non-wetting conditions and exhibit initially spherical shapes [5]. We then add 1.24 µM G-actin to the system, leading to rapid uptake of G-actin and subsequent in-droplet polymerization of F-actin visualized using fluorescently labelled phalloidin. We allow for continued in-droplet polymerization and record the shapes using a confocal microscope about 13 h after addition of actin (Materials and Methods).

Exemplary shapes are shown in Fig. 1. We find that filaments that exceed the diameter of the condensate localize at the condensate interface to minimize their bending energy as previously described [12]. In large condensates, this results in F-actin decorating the interface, see Fig. 1(a). In smaller condensates, filaments accumulate progressively in one ring, Fig. 1(b). In yet smaller condensates, the filament ring contributes more strongly to the emerging shape, resulting in oblate spheroids (elsewhere sometimes referred to as “discs”), see Fig. 1(c). Recording the aspect ratio (AR), i.e. the ratio of major to minor principal axis of an ellipsoid fitted to each shape, we find that high aspect ratios occur predominantly in condensates of smaller volume, Fig. 1(d). This is in line with the consideration that filaments have to comply to smaller radii in smaller condensates, leading to a higher contribution of the bending energy and increasing deviations from the initially spherical shape preferred by minimizing the surface energy between condensate and buffer.

**FIG. 1.**
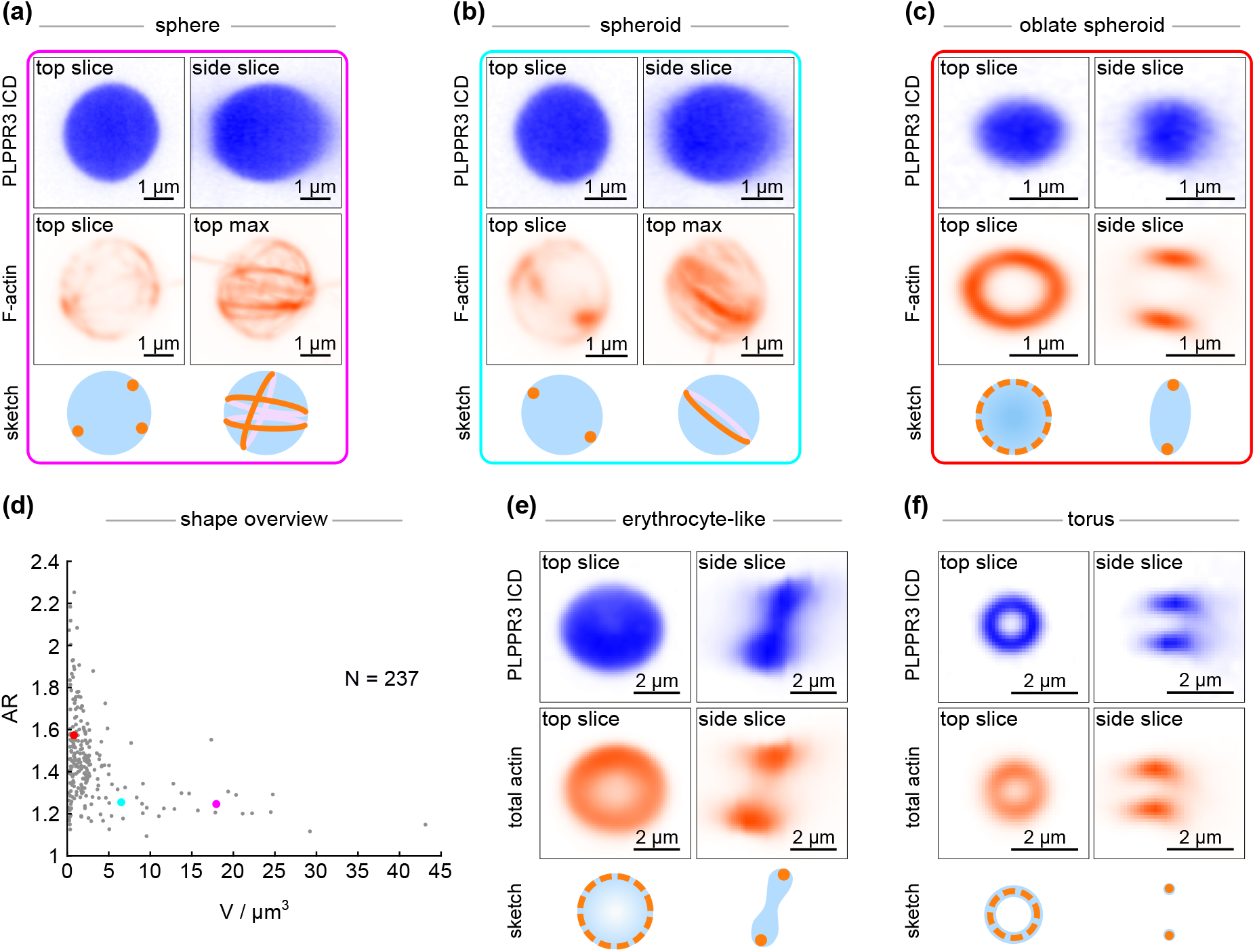
Polymerization of F-actin inside biomolecular condensates induces mutual shape deformations of condensates and embedded filaments. (a)-(d) Confocal images of condensate-filament shapes from an experiment containing 20 µM PLPPR3-ICD and 1.24 µM G-actin. (d) Aspect ratio as a function of droplet volume for N=237 condensates from the same experiment. The condensates from panel (a), (b) and (c) are highlighted in magenta, cyan and red respectively. (e)-(f) Condensate-filament shapes recorded at a higher concentration of actin (4 µM) and lower concentration of PLPPR3 ICD (10 µM) show more pronounced deformations, resembling an erythrocyte shape (e) or torus shapes (f). The aspect ratio overview for this dataset is shown in Supplementary Fig. S1.

Systems with higher actin and lower condensate concentration can reach more extreme shapes. Increasing the actin concentration to 4 µM and decreasing the PLPPR3 concentration to 10 µM results in shapes with higher aspect ratios. In this system, we observe new morphologies similar to the shape of erythrocytes, where the condensate phase forms an increasingly thin film towards the center of the actin ring, see Fig. 1(e). Condensates, often the smallest in the respective experiment, may further form torus shapes, where condensate and filament indiscernibly localize in one ring, Fig. 1(f).

These experimental results reproduce previous observations of biomolecular condensates that fully containt F-actin filaments [12–14, 17]. They suggest a competition of bending energy of the filament and surface energy of the condensate against the buffer that underlies the observed shape changes, in line with the assumptions in other studies [13, 18].

### B. Analytical Model

Inspired by the experimental results, we construct an analytical model on the assumption that surface and bending energies dominate the shapes of droplets containing entire filaments. The total energy is then given by

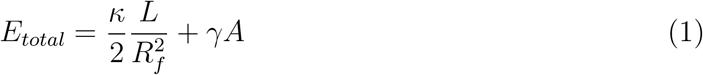

with bending stiffness of the filament *κ*, filament length *L*, bending radius of the filament *R*_*f*_, surface tension *γ* between droplet and environment, and surface area of the droplet *A*. The energy is thus defined by three geometric parameters, *L, R*_*f*_ and *A*, and two material properties, *κ* and *γ*. We approximate the shapes observed throughout the experiments as solids of rotation, from initial sphere, to oblate spheroid, modeled as a lens, to torus. These geometries link *V, A* and *R*_*f*_ to a defined shape, see Materials and Methods. The spatial extent of the filament is assumed to approach zero such that it can overlap, and the filament is localized at the largest radius of the droplet in each geometry.

Fig. 2(a) shows an exemplary energy landscape for a set of *γ, κ, V* and *L* upon varying the filament radius *R*_*f*_ for two geometries to determine the minimal energy shape. The material parameters *κ* and *γ* are set to 2 · 10^−25^ Jm and 10^−5^ J/m^2^ respectively, which are plausible values for F-actin and biomolecular condensate *in vitro* [22, 23]. Varying the filament radius *R*_*f*_ changes the droplet geometry and therefore *E*_*total*_, allowing for the determination of the minimal energy shape. In this way, the model can predict the minimal energy shape for any set of *V, L, κ*, and *γ*, as long as the predefined shapes are geometrically accessible. Fig. 2(b) shows the resulting shape phase diagram. The model predicts increasingly oblate shapes upon increasing *L* or decreasing *V*. Moreover, it predicts a transition from spheroid to torus shapes, particularly as volumes decrease. The analytical model thus recapitulates the shape changes observed throughout in-droplet polymerization of F-actin in PLPPR3 ICD condensates (Fig. 1).

**FIG. 2.**
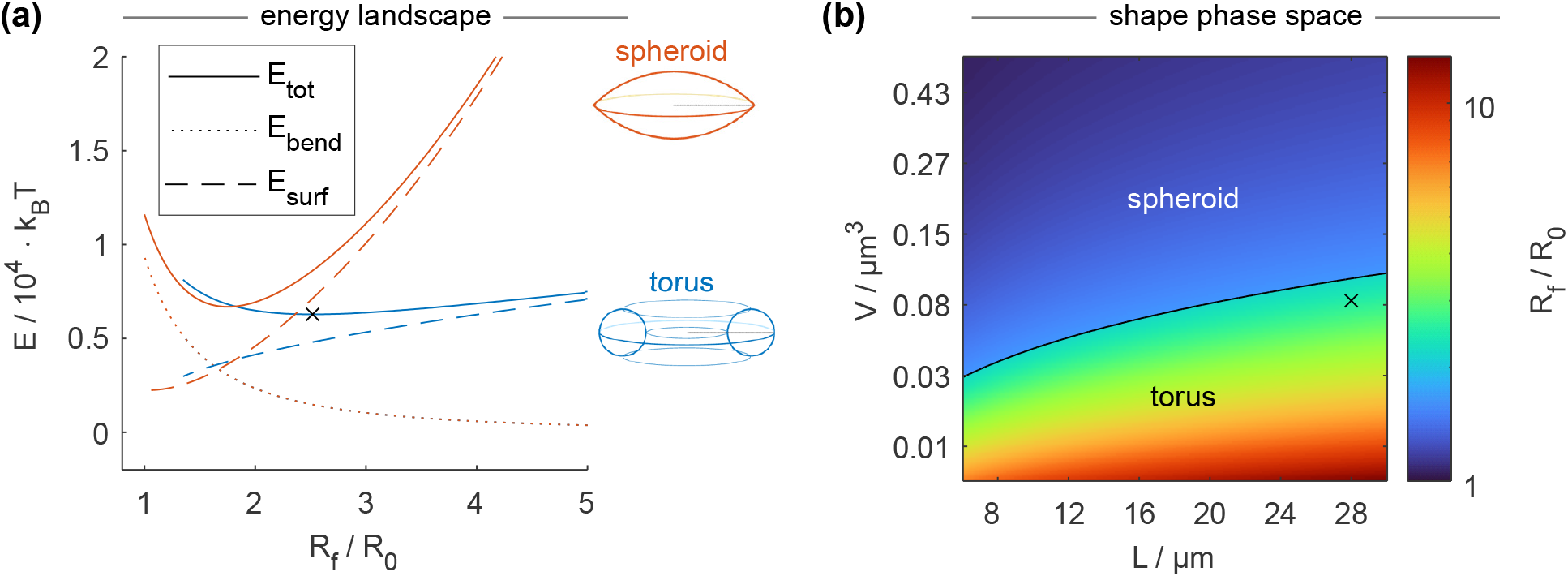
The analytical model predicts shape deformations as a function of bending rigidity *κ*, surface tension *γ*, droplet volume *V* and filament length *L*. (a) Energy landscape for a droplet with fixed material properties, constant volume *V* and filament length *L* upon varying the geometric shapes. The shape is determined by the variable *R*_*f*_, which is the maximal radius the filament may assume in a given shape, indicated by the dashed line in the sketches of the spheroid and torus to the right. The filament radius *R*_*f*_ is normalized by *R*_0_, which is the radius of a sphere with the same volume. The black x defines the minimal energy shape, in this case a torus. (b) A shape diagram of the minimal energy shape for varying volume *V* and filament length *L*. The surface tension *γ* and bending rigidity *κ* are fixed at *γ* = 1 · 10^−5^ J/m^2^ and *κ* = 2 · 10^−25^ Jm.

The analytical model is free of fit parameters, allowing for quantitative comparison to experimental data. However, the small size and experimental constraints of the PLPPR3 ICD condensates containing F-actin limit the quantitative analysis of the emerging shapes, e.g. in the determination of *L*. The model should be applicable as long as surface and bending energies dominate the energy landscape. We expect an upper limit for the applicability as soon as contributions of gravity become apparent, i.e. as the droplet approaches the capillary length. To probe the applicability of the model at larger and more accessible scale, we place a water droplet below but near the capillary length (*V* ≈ 9 µL) onto a super hydrophobic substrate and embedded a thin polyester filament (polylactic acid (PLA), 26 µm diameter, 24.788 mm long) into the droplet. Supplementary Fig. S2(a) and (b) show exemplary shapes and the AR of the water-polyester system as the water droplet evaporates. This experiment shows an increase in the aspect ratio as the volume decreases, in line with the observations from condensates with F-actin and the analytical model. We extract the volume *V* from the cross-sectional area of the droplet and determine the length and diameter of the filament on a microscope after the experiment. Assuming *γ* = 72 mJ/m^2^ as the surface tension of water against air under ambient conditions and using our measured droplet volume and polyester filament length, we can utilize the analytical model to predict the bending stiffness *κ* of the polyester filament. We do so by matching the observed droplet shapes from the experiment to the predicted shapes of the analytical model. The result is shown in Supplementary Fig. S2(b) and suggests *κ* ≈ 1.5 · 10^−11^ Jm. Approximating *κ* via the second moment area *I* (calculated from the filament diameter) and Young’s Modulus *E*_*y*_ (given by supplier) as *κ* = *E*_*y*_*I* suggests *κ* in the range of 5 to 9 · 10^−11^ Jm. Considering experimental uncertainties, especially effects of wetting on the substrate, the analytical model accurately captures *κ*, showing quantitative agreement between experiment and analytical model.

While the water-polyester system provides quantitative agreement, we can not generalize these findings to biomolecular condensates due to different scalings of filament length and liquid volume. The ratio of circumference to volume for a sphere scales with the inverse squared radius, i.e. increases for smaller volumes. The volume of biomolecular condensates is about nine orders of magnitude smaller than the volume of the water-polyester system. Although the analytical model (Fig. 2) suggest a similar amount of filament loops in condensates compared to the water-polyester system, the ratio of droplet volume to filament length increases as the droplet volume decreases, suggesting more pronounced contributions of the interactions between filament and liquid phase in condensates.

### C. Simulations

To investigate the physical mechanisms of mutual deformations between biomolecular condensates and contained cytoskeletal filaments at their native length and volume scale, we perform coarse-grained molecular dynamics simulations using LAMMPS [24–30] that resemble biomolecular condensates interacting with F-actin. The simulation contains two particle types representing the fluid phase (liquid droplet and surrounding gas) and the filament. Particles of the fluid phase interact with each other via a truncated and shifted Lennard-Jones potential, allowing for spontaneous phase separation. The depth of the potential well *ϵ*_*ll*_ = 1*/*0.65 k_*B*_T and particle size *σ* = 0.03 µm result in an effective surface tension of the liquid against gas phase of 3 · 10^−6^ J/m^2^ (see Materials and Methods). This matches the order of magnitude expected for biomolecular condensates *in vivo* [22, 31]. The number of particles is chosen to achieve volumes comparable to biomolecular condensates *in vivo*, with larger simulated droplets forming radii of up to 0.4 µm.

The filament is modeled as a chain of self-avoiding beads. The bending rigidity and stretchability of the filament are implemented via (angular) harmonic springs with spring constant *κ*_*b*_ = 3.34 · 10^−18^ Nm and *κ*_*s*_ = 0.17 N/m, respectively, set to resemble F-actin (see Materials and Methods). The interactions between liquid and filament particles are regulated via another truncated and shifted Lennard-Jones potential with the same *σ* as used for the interactions between fluid particles, but with a slightly deeper potential of *ϵ*_*lf*_ = 1*/*0.6 k_*B*_T, ensuring high effective wettability and incorporation of the filament into the droplet. The simulations are performed in a 2 × 2 × 2 *µm*^3^ simulation box with periodic boundary conditions and evolve according to Langevin dynamics in the NVT (canonical) ensemble at *T* = 300 *K*. For the Langevin dynamics, the damping factor, particle mass, and time steps, are chosen to achieve an effective viscosity resembling that of the water at around 1 mPas, see Materials and Methods.

#### 1. Non-self-avoiding filaments simulations

First, we perform simulations resembling the assumptions of the analytical model and water-PLA system but at the volume scale of biomolecular condensates. In particular, filament loops may overlap such that all loops assume *R*_*f*_ of a given shape, resulting in filaments that take up comparably little volume. This is implemented by setting the interaction potential between non-sequential filament particles to zero. Throughout these simulations, the filament length is fixed at 400 *σ*, which corresponds to about 12 µm. We then simulate the system with different initial conditions, varying the number of liquid particles to change the volume of the droplet and the initial configuration of the filament to change its radius of curvature *R*_*f*_. Note that the filament bundle quickly becomes kinetically trapped and does not freely rearrange in the simulation, whereas the fluid phase can.

After relaxation of the system, we can observe two different topologies, spheroids (contractible topology) and structures with a discontinuous liquid phase in the center (non-contractible topology), see Fig. 3(a). For the later, we observe two different shapes, a torus shape of (near) constant diameter of the liquid along the azimuthal angle *ϕ* and a kettlebell shape with varying liquid diameter along *ϕ* (Supplementary Fig. S3). We utilize the distribution of liquid along *ϕ* to differentiate torus and kettlebell, see Materials and Methods. Throughout the simulations, we find spheroids for larger volumes and smaller *R*_*f*_. Tori occur at the lowest volumes, and kettlebell shapes form between the spheroid and torus regimes, covering an increasing area of the shape space as *R*_*f*_ increases.

**FIG. 3.**
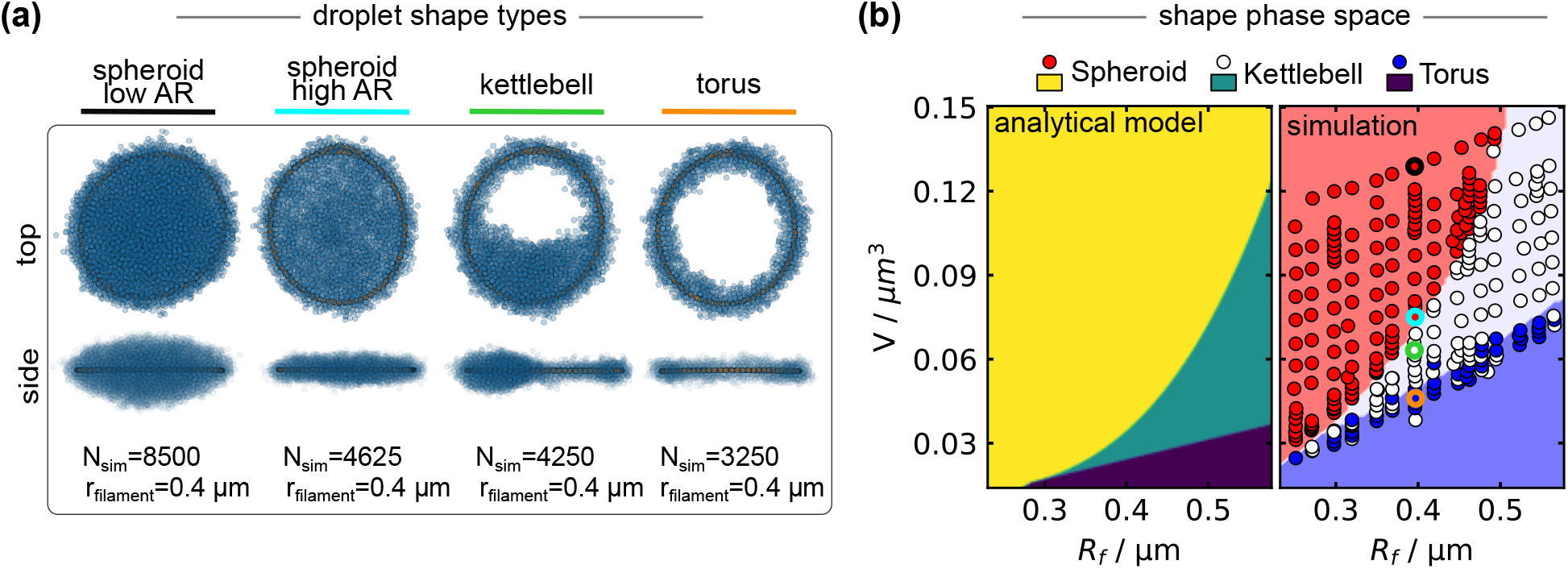
Simulations with overlapping filaments reveal an additional geometry and align with an extended analytical model. (a) Top and side view of exemplary shapes resulting from the simulation. The shown shapes have identical filament length and *R*_*f*_ and differ only in *V*. Filament particles are depicted in orange, fluid particles are depicted in slightly transparent blue. Filament loops lie mainly on top of each other. (b) Simulation results (right) compared to the shape diagram determined via the analytical model for *L* = 12 µm with varying *V* and *R*_*f*_. The points corresponding to the simulation results in (a) are highlighted in their respective color. The background color is predicted by a support vector machine trained on the simulation data (see Material and Methods).

Taking a closer look at the organization of the liquid particles around the filament in kettlebell and torus shapes, we find hexagonal packing of the liquid particles around the filament. We attribute this to a high wetting energy, amplified by the overlapping filaments, which results in a pronounced wetting film, see Supplementary Fig. S4(a) and (b). We observe that the transition from torus to kettlebell occurs when the liquid volume exceeds the volume retained within this wetting film, see Supplementary Fig. S4(c). The remaining liquid then arranges in shapes that minimize the surface energy of the system, resulting in the kettlebell geometry. Comparable shape instabilities, similar to the Rayleigh-Plateau instability but resulting in a single liquid bulge, have been observed in other wetting experiments [32, 33].

In this way, the kettlebell geometry is in line with the assumption of the analytical model that surface and bending energy define minimal energy droplet shapes, but it requires additional constraints in order to account for the observed wetting effects. To incorporate the kettlebell into the model, we approximate the kettlebell shape as a torus surrounded by a finite liquid film with an additional sphere formed by the excess liquid. The maximum film thickness around the filament affects the availability of the torus and kettlebell shape and needs to be known a-priori or fitted. Setting the film thickness to match the thickness observed in the simulations, we can compare the analytical model against the simulation results. Fig. 3(b) shows the analytical prediction for material properties matching the simulation conditions with fixed filament length *L*, and varying *V* and *R*_*f*_. We find that the different shapes predicted by the analytical model after introducing the kettlebell occupy distinct areas inside the diagram, that align well with the simulation results in their relative position. Moreover, the model places the shape regions in the correct order of magnitude of the geometric parameters. However, shape transitions consistently occur at higher volumes in the simulation compared to the analytical prediction, leading to a misalignment between simulation and model. This discrepancy is likely caused by the infinitely small volume of the filament and lack of any wetting film in the analytical model.

Note that implementing a maximal film thickness as a shape constraint and the addition of the kettlebell lead to an expansion of the regime in which the spheroid is favorable as fewer shapes are available in the torus regime. The emerging kettlebell regime, does not make up for this effect, see Supplementary Fig. S3(a) and (b).

Overall, the coarse-grained molecular dynamic simulations at the volume scale of condensates qualitatively reproduce the trends observed in the experimental findings. However, the simulations reveal kettlebell shapes as an additional morphology that we do not observe in experiments. Implementing the kettlebell shape in the analytical model by prescribing a maximal film thickness, on the other hand, matches simulation results onto the analytical model. This shows that the initial assumption that surface and bending energies define minimal energy shapes still holds. However, the emergence of the kettlebell shape points towards wetting effects as additional contributors that affect the shapes of droplets containing filaments.

#### 2. Self-avoiding filament simulations

To more accurately capture wetting effects of droplets and contained filaments on the scale of biomolecular condensates, we next model the filament as non-overlapping. This ensures more realistic scaling of the interfacial area between filament and liquid and consideration of the filament volume. We implement the self-avoiding filament by using only the repelling part of the Lennard-Jones potential for non-sequential filament beads. In contrast to the simulation with overlapping filaments, the filaments are free to rearrange, enabling the system to relax into minimal energy shapes by adjusting both liquid volume and filament organization. Changing *V* and *L* in the simulation by altering the number of liquid or filament particles, we can explore the space of minimal energy shapes.

Fig. 4(a) shows the types of the shapes we observe in the simulation after relaxation. These are the same shapes as in the simulation with overlapping filament, with spheroids of various aspect ratios, torus and kettlebell shapes. The top and side views of the representative shapes in Fig. 4(a) show that, contrary to our first model and in agreement with our assumption, the filament occupies a relative large fraction of the shape volume (between 5% and 20%). While filaments localize to the edge of the given shape, they must now also occupy space further from the interface due to the self-avoidance. Filament loops are separated from each other by a thin layer of liquid particles, forming a filament bundle at the edge of the respective shape that also contains a significant amount of liquid.

**FIG. 4.**
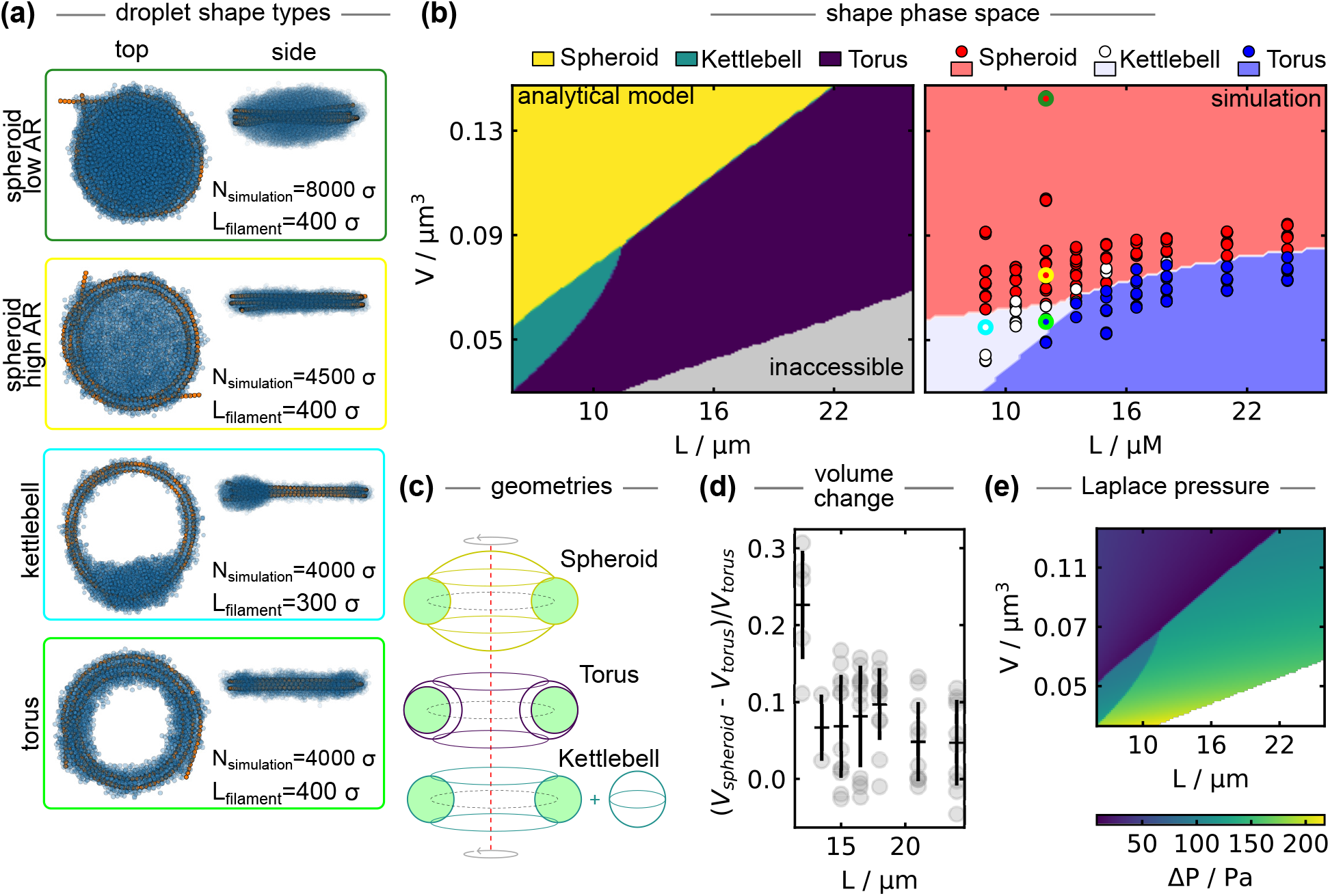
Simulations of self-avoiding filaments reveal minimal energy shapes and provide insights into shape transitions. (a) Top and side views of emerging shapes after equilibration. (b) Shape space predicted by the modified analytical model (left) compared to simulation results (right). The background color for the plot of the simulations is predicted by a support vector machine trained on the simulation data (see Material and Methods). (c) The modified analytical model considers an extended filament bundle (green torus) containing the filament and a fraction of the liquid. This filament torus is augmented with two spherical caps for the spheroid shape, surrounded by a larger torus of liquid for the torus shape, accompanied by an additional liquid droplet for the kettlebell shape. (d) Loss of volume upon transitioning from spheroid to torus topology as a function of filament length. (e) Estimated Laplace pressure throughout the shape diagram of the analytical model.

To adjust our analytical model to this finding, we replace our previously dimensionless (infinitely thin) filament with a filament bundle in the form of a torus, inside which the filament takes up only a fraction *ε* of the volume. The remaining volume is occupied by the liquid phase. We set *ε* = 0.27, which aligns with geometrical considerations, see Materials and Methods. The filament length *L* and the volume fraction of the filament *ε* therefore define the volume of the filament bundle. In case of the torus, the (excess) liquid phase is now organized around this filament bundle, whereas it forms a separate sphere in kettlebell shape. This is different to our previous analytical model as there is prescribed film thickness anymore. The spheroid shape is modeled by adding spherical caps to the top and bottom of the generating circle of the filament bundle, see Fig. 4(c).

Fig. 4(b) shows the predictions of the modified analytical model next to the shapes observed in simulations. Compared to the simulations with overlapping filaments and our analytical model with a dimensionless filament, we find that the kettlebell shapes now occur on the opposite side, at small *V* and small *L*, both in simulation and analytical model. We attribute this behavior to liquid captured as a wetting film inside the filament bundle. As the filament becomes shorter, less liquid is captured in the bundle, allowing for the remaining liquid to arrange in shapes that minimize surface area, analogously to the emergence of the kettlebell shape before. In this way, the shift of the kettlebell regime is a consequence of the more realistic consideration of wetting conditions. Interestingly, the asymmetry of the liquid phase observed in the kettlebell shape appears to be present already at the onset of the shape transition from a spheroid, see Supplementary Fig. S5.

Similar to the comparison between analytical model and simulations with overlapping filaments, we find that the modified analytical model captures the qualitative phase behavior but the exact extent of the different regimes is miss-aligned. This is likely caused by the simplified geometries and assumptions in the model that only estimate the actual shapes, e.g. the simple geometry of the filament bundle torus. Moreover, the model assumes a fixed ratio of liquid in the filament bundle, which may realistically change upon differences in bending energy or available liquid. The qualitative but limited quantitative agreement, substantiates the conclusion that surface tension and bending rigidity still present the driving forces for shape changes of condensates containing entire filaments, albeit within constraints set by the wetting interactions between liquid and filament.

In addition to the steady-state behavior, the simulations provide insights into the dynamics of the shape relaxation process that neither the analytical model nor our experimental data provide. Supplementary Fig. S5(c) shows how *E*_*bend*_, *E*_*surf*_ and total *E*_*total*_ change throughout the shape relaxation of a representative simulation. We find that *E*_*bend*_ is the larger contributor to *E*_*total*_, driving the energy minimization. Decreasing *E*_*bend*_ comes at the expense of creating additional surface area between fluid and gas phase, resulting in increasing *E*_*surf*_, however, the total energy of the system still decreases. While *E*_*surf*_ relaxes continuously, we observe that changes in *E*_*bend*_ may occur more abruptly, indicating that the energy landscape of *E*_*bend*_ is shallow (particularly towards larger *R*_*f*_) and that the relaxation process may be hindered, e.g. by entanglement of the filament loops. This matches with the observation that simulations of the same condition do not always result in the same *E*_*total*_ and corresponding shape, see Supplementary Fig. S5(d). A similar behavior can be observed in the experiment with the water-polyester system where the semi-major axis of the shape, defined by the filament, shows abrupt changes, indicating the presence of static friction and/or filament entanglement also in experiments, see Supplementary Fig. S2(d). While the analytical model predicts the minimal energy shape, effects of static friction and entanglement may consequently delay or hinder the relaxation to this exact shape in experiments and simulation.

We further find that changes in topology lead to changes in liquid volume. The volume of the liquid phase only changes minimally throughout shape relaxations that do not change topology. However, we observe a pronounced reduction of liquid volume upon the transition from spheroid to torus topology, see Fig. 4(e) and Supplementary Fig. S5(b). The observed loss in volume is smaller for longer filaments. We attribute this behavior to the wetting film around the filament, that effectively traps an increasing fraction of the liquid as the filament length increases. The shape transition from spheroid to torus is accompanied by an increase in the curvature of the fluid-gas interface, suggesting an increase in Laplace pressure of the fluid phase. Excess liquid not stabilized by favorable wetting interactions may transition to the gas phase due to the increasing Laplace pressure. Estimating the Laplace pressure of a given shape in the analytical model by the curvature of the liquid phase, omitting the wetting-stabilized filament bundle, we find a pronounced increase in pressure upon transitioning from spheroid into kettlebell or torus, see Fig. 4 (e). Moreover, the Laplace pressure drops throughout shape changes towards increasingly oblate spheroids due to the flattening spherical caps. Contrary to shape changes of the spheroid, increasingly oblate torus shapes lead to a further increase in Laplace pressure. We anticipate that increasing (decreasing) Laplace pressure leads to efflux (influx) from the liquid to the gas phase. This is particularly clear in the loss of volume upon shape transition from spheroid to torus in the simulation. As soon as multiple droplet-filament shapes of varying size and geometry are present, as in our experiments, changes in Laplace pressure are expected to lead to Ostwald ripening. This suggests that particularly torus and kettlebell shapesare susceptible to loss of volume through Ostwald ripening until all liquid not captured in the filament bundle disappears. This may explain why we do not observe kettlebell shapes in experiments with condensates and F-actin and why the liquid and filament phase perfectly overlap in experimentally observed torus shapes. Conversely, shape changes towards increasingly oblate spheroid shapes may lead to decreasing Laplace pressure, stabilizing such oblate shapes.

## III. DISCUSSION

Liquid droplets containing entire filaments show characteristic mutual deformations. We find that emerging shapes are governed by a competition of bending and surface energies across length scales. However, as the ratio of filament length to liquid volume increases, wetting interactions set increasingly noticeable boundaries within which surface and bending energy define minimal energy shapes.

Throughout simulations, we implemented a higher affinity between fluid particle and filament than between fluid particles themselves, leading to prewetting film formation on the filament. We anticipate that the relative contribution of wetting effects on the emerging shapes of filament-containing droplets decreases as the affinity for the filament decreases, leading to less liquid trapped around the filament. However, wetting effects should continue to contribute to the emerging shapes as long as the droplet has a higher affinity for the filament than for the gas phase. Adhesion of the filament to the droplet may further be aided by interactions with the interface through Pickering-like effects [3].

Our work expands upon the classical stability consideration of the bendocapillary length [15], which defines the radius at which the adhesion of a filament to a spherical droplet equals the bending energy necessary for the filament to conform to the spherical droplet shape. We show that droplets may retain filaments also at sizes smaller than the bendocapillary length through mutual shape deformations. This will be the case if an initially stable shape of a droplet containing a filament (e.g. forming above the bendocapillary length or by polymerizing filaments inside the droplet) is modified by extracting volume (due to evaporation or ripening) or by elongating the filament through polymerization. However, the resulting stable minimal energy shape below the bendocapillary length is likely not accessible by mere contact between a spherical droplet of the same volume, now below the bendocapillary length, and a filament of the same length. Consequently, shape deformations expand the regime of accessible shapes but also introduce hysteresis. Moreover, while the bendocapillary length is independent of filament length, the emerging shape deformations are not.

Fig. 5 (a) exemplifies this effect for a condensate with plausible surface tension *in vivo* interacting with a 20 µm long filament with varying bending rigidity, here expressed as the filament’s persistence length. The shape space is calculated using the fit-free analytical model (neglecting wetting effects and with dimensionless filament). The red crosses show an estimate of the bendocapillary length for selected biopolymers. The white lines indicate shapes below the bendocapillary length, in which the bending energy of the filament is below the adhesion energy assumed in the calculation of the respective bendocapillary length (see Materials and Methods). We find that mutual shape deformations enable smaller liquid volumes to capture different biopolymers. While we find negligible increase for interactions with microtubules, shape deformations markedly extent accessible shapes for condensates interacting with comparably more flexible F-actin. Particularly, torus shapes emerge as a generally stable geometry. Strikingly, shape deformations make all shapes accessible for softer biopolymers such as intermediate filaments (IFs) or DNA, even only considering spheroid shapes (Supplementary Fig. S6). This not only shows that mutual shape deformations expand accessible shapes, but also that filaments, once captured in a droplet, can remain entrapped in the droplet phase even as the volume decreases or filament length increases.

**FIG. 5.**
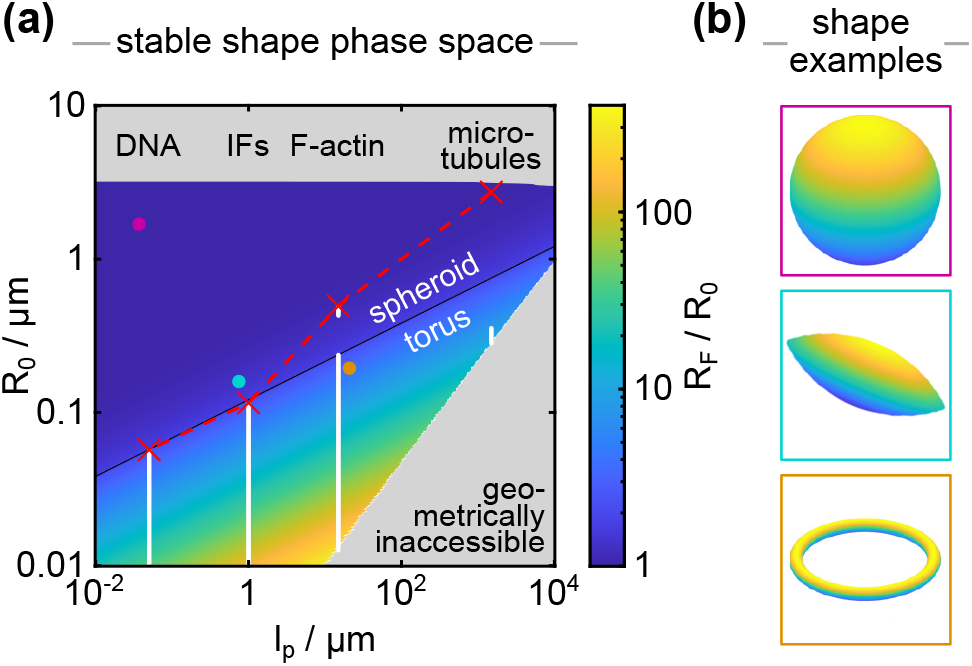
Mutual deformations of biomolecular condensates and different biopolymers expand the regime in which droplets retain the filament(s). (a) Shape diagram of minimal energy shapes for a condensate with a surface tension of 5 · 10^−6^ J/m^2^ and filament length of 20 µm as a function of condensate volume and filament bending rigidity. The bending rigidity is expressed in terms of persistence length and the filament volume in terms of radius of a sphere of the same volume *R*_0_ for convenience. The colormap shows the aspect ratio of the minimal energy shapes as *R*_*f*_ */R*_0_. The red crosses indicate the approximated *R*_*bc*_ for microtubules, F-actin, intermediate filaments and DNA. All shapes above this line are stable. The white dots indicate additional shapes stabilized by mutual deformations. (b) Examples of predicted shapes as marked by the colored dots in panel (a).

Further, we identify changes in liquid volume upon shape and, particularly, topological changes, which may carry functional significance. Chemically active droplets have been suggested as early precursor structures of life [34]. In this context, polymerization of filaments within such droplets would lead to shape changes and an increase in surface area, possibly favoring uptake of metabolites. Moreover, increasingly oblate spheroid shapes could facilitate droplet growth through Ostwald ripening. Emerging shapes of droplets with filament rings further strongly resemble structures formed by actin networks polymerizing within lipid vesicles, a minimal model for membrane-enclosed life [35].

Aside from canonical biopolymers, filamentous structures may also form due to abarrant protein interactions. Amyloids are highly ordered protein aggregates that form rigid, filament-like structures that are implicated in various diseases [36]. These structures may also form within biomolecular condensates, particularly at the interface [37]. Engineered condensates may be capable of retaining such filaments, and the resulting shape deformations could serve as indicators of progressing amyloid formation.

Combined, these findings show that droplets and filaments have more complex and detailed combined behavior than previously recognized. In the context of biomolecular condensates, our results highlight their capacity for potent filament entrapment and (re)structuring and provide a physical framework for more detailed understanding of such interactions.

## ACKNOWLEDGMENTS

We thank Dr. Nils Köster and the team of the Berlin Botanic Garden and Botanical Museum for supplying the taro (*Colocasia esculenta*) plant used as a substrate in experiments with water and PLA filament. Imaging of the experiments with PLPPR3 ICD and actin was done at the Advanced Medical BIOimaging Core Facility of the Charité-Universitätsmedizin Berlin (AMBIO). TJB acknowledges financial support from EMBO Postdoctoral Fellowship ALTF 625-2022. HK acknowledges funding from UKRI EPSRC (grant EP/V034154/2). HK and SG acknowledge funding from Research Council of Norway (grant no. 335901).

TJB developed the initial project idea, which has been realized and expanded with all authors. TJB, SB performed experiments. SB purified proteins. TJB, FW analyzed experiments. FW, HK performed and analyzed simulations. TJB, FW developed and analyzed the analytical model. BJE, RLK, SR, SNG and HK provided resources. FW and TJB wrote the paper with input from all authors.

The authors declare no competing interests.

## Appendix A

**Materials and Methods**

### 1 Experiments

#### a. Actin-protein condensate experiments

PLPPR3 ICD was purified as described in [5]. Experiments are similar to actin polymerization assays described in [5]. 20 µM PLPPR3 ICD (3% fluorescently labeled using NHS-ester conjugated to Alexa-488) was incubated in 20 mM HEPES (pH 6) with 150 mM NaCl, 5 mM DTT and 200 nM phalloidin-Atto 565 (Merck). The experimental volume further included 1:10 F-actin buffer (10x stock consisting of 1 M KCl, 20 mM MgCl_2_, 0.1 M imidazole, and 10 mM ATP at pH 7.4). The experimental volume was pipetted into glassbottom dishes coated with polyvinyl alcohol (PVA, #363065, Sigma) to achieve non-wetting conditions of the condensate. Droplets were allowed to form and settle for 2 h before adding 1.24 µM G-actin (31% fluorescently labeled, both from Hypermol, #8101-01 and #8158-01). All concentrations were calculated for a final volume of 4 µL after adding actin. Actin selectively partitions into the condensates and polymerizes within a lag time of about 30 min [5]. Images were taken on a Ti2 Nikon Eclipse confocal microscope with 60x oil objective and 4x SoRa mode with a step size of 0.1 µm in z after overnight incubation. F-actin was visualized using the phalloidin stain.

The high actin concentration, low PLPPR3 ICD concentration experiments were performed by first forming F-actin in the same buffer as above but with 4 µM actin and without phalloidin, before adding PLPPR3 ICD. 10 µM PLPPR3 ICD was added and images were taken 30 min after addition of PLPPR3 ICD on a Ti2 Nikon Eclipse confocal microscope in dual camera mode with 60x oil objective with a step size of 0.3 µm in z. The addition of PLPPR3 ICD lead to rapid formation of condenstes, leading to rapid uptake of actin into the condensate. Actin was visualized using its fluorescent tag, i.e. showing both G- and F-actin.

Both datasets were analyzed using Matlab. First confocal stacks were rescaled to achieve isometric voxel size using the matlab function imresize. Note that this step did not involve deconvolution and therefore retains the point-spread-function of the optical setup with lower resolution in the z-direction. Condensates were detected in the PLPPR3 channel using a heuristic threshold based on 60% intensity between background and maximum intensity (99.9 percentile) inside each condensate. The F-actin channel visualized via phalloidin staining was thresholded similarly as 75% intensity between background and the maximum F-actin signal in the condensate (99.9 percentile). Both image masks were combined to determine the principal axis (PA) ratio using the matlab function pregionprops3.Droplets that exceed the imaged volume were not considered, leading to a cut-off at larger condensate volumes. Note that the non-isometric point spread function results in over-estimation of droplet shapes in the z-direction.

#### b. PLA filament-water droplet experiments

Droplets of MilliQ distilled water were placed on leaves of *Colocasia esculenta*, more commonly known as taro, which exhibit super-hydrophobic properties. Leaves were gently mounted onto an xyz-micromanipulator and imaged on a horizontal microscope composed of a Lensagon TC5M-10-65 telecentric lens with Imaging Source DFK 33UX249 camera. Filaments of polylactic acid polyester (PLA, 3D printing filament, Orbi-Tech, Germany, *E*_*y*_ ≈ 3.54 MPa) were pulled from a heated filament at close to 200°C to achieve thin filament diameters on the range of tens of micrometers. The filaments were manually inserted into the condensate using tweezers and plasma-cleaned prior to experiments to improve wettability. After timelapse imaging, the filament was mounted onto a coverslip to determine the diameter and length on an upright microscope.

### 2. Analytical model

#### a. Geometries

In the analytical model, we fix the volume of a given droplet and calculate the minimal energy shape for filaments of varying length. The reference volume is a sphere of radius *R*_0_ with volume 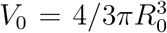. The lens and torus shapes are generated as solids of rotation, see Fig. A1.

The lens shape is generated by rotating a circular segment along the normal through its apex, see Fig. A1 (a). This geometric shape can also be interpreted as two identical but flipped spherical caps joined at their circular base. The lens can be characterized with two parameters, the height of one spherical cap *h* and the radius of the circular base *a*.

We then find the surface area as

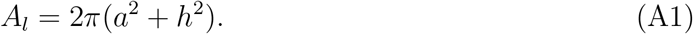

The volume is given as

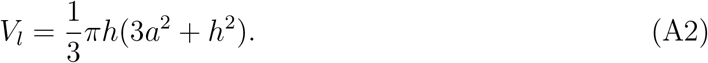

Fixing the volume to the volume of the original, spherical droplet, we find

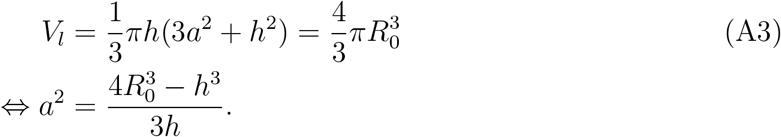

**FIG. A1.**
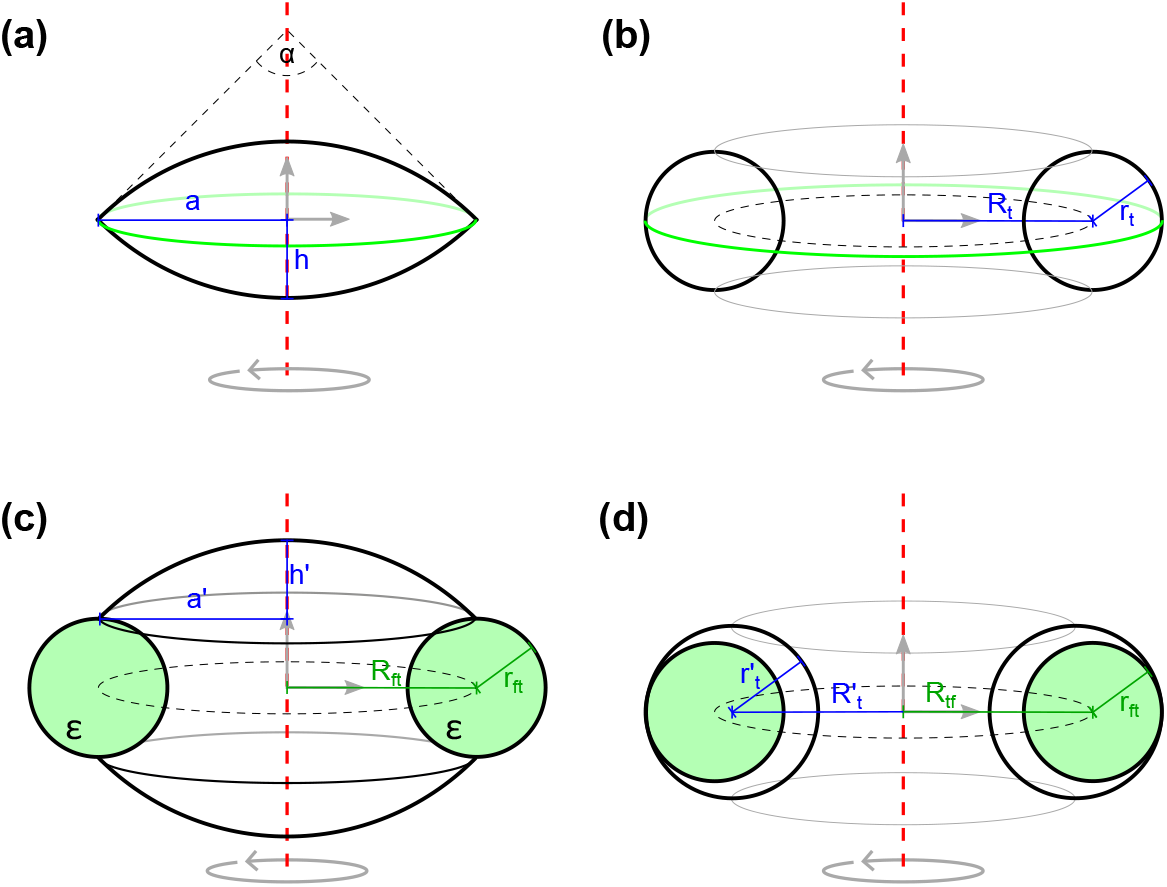
The lens and torus shape in the analytical model are solids of rotation. (a) The lens is generated by rotating a circular segment (black line) around an axis of rotation (red dashed line). (b) The torus is generated by rotating a full circle (black line) around an axis of rotation (red dashed line). (c) Modification of the lens shape upon adding a filament bundle. The filament bundle has a radius of rotation *R*_*ft*_ and a radius of the generating circle *r*_*ft*_. Two spherical caps are added on the top and bottom of the generating circle relative to the axis of rotation. Note, that the lens can also be concave for a shape similar to erythrocytes. The filament bundle has a volume fraction of embedded filament of *ε*, the remaining volume is taken up by the liquid phase. (d) Modified torus shape engulfing a filament bundle with geometric constraint 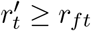.

The surface area is then

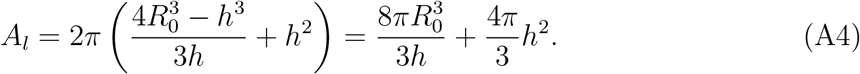

The maximal radius the filament can assume is

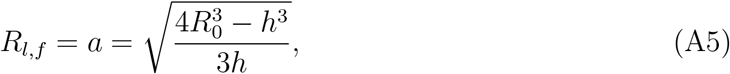

which is well defined as *h* ∈ (0 *R*_0_] and *a >* 0. But *A*_*l*_ and *R*_*l*_ are functions of *R*_0_ and *h*. With a fixed *R*_0_, we can thus describe different volume conserving lens shapes by varying *h*.

The torus is generated by rotating a circle of radius *r*_*t*_ at a distance of *R*_*t*_ from the origin, see Fig. A1 (b). Applying Pappu-Guldinus theorems we find the torus surface area is

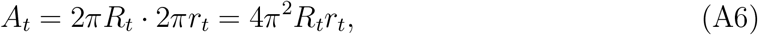

and for volume is

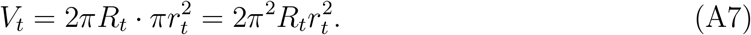

Fixing the volume of the torus to the volume of the reference sphere, 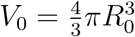, we find

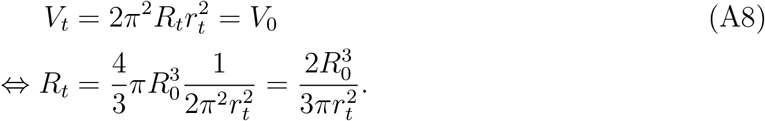

For the surface area, we thus find

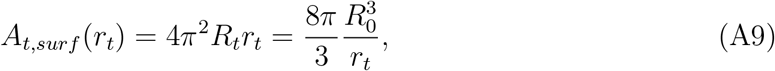

i.e. we minimize the surface area if we maximize *r*_*t*_. In the torus shape, a filament can assume the maximal radius

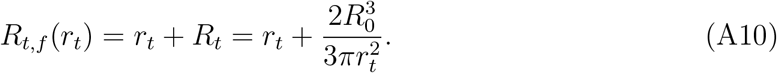

Varying *r*_*t*_ thus allows to calculate all plausible *R*_*t,max*_ and *A*_*t,surf*_ for a fixed *R*_0_.

To consider an extended bundle of filaments, we introduce a torus with radius of rotation *R*_*ft*_ and radius of the generating circle *r*_*ft*_ into the lens and torus shapes. This filament torus is composed of both filament and liquid phase, with a filament volume fraction *ε. ε* is estimated from hexagonal packing of filaments with an additional wetting film of the liquid surrounding each filament, leading to a larger distance between filaments, see below.

We model the lens shape with engulfed filament bundle as the filament torus with added spherical caps at the top and bottom of the generating circle of the filament torus, see Fig. A1 (c). The volume of this shape is composed of a cylinder with radius 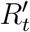 and height 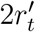, a “half-torus”, i.e. the volume of the torus that extends beyond 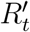, and the volume of the two spherical caps. The volume of the “half-torus” can be calculated using Guldinus first theorem by rotating a half-sphere around the radius between the axis of rotation and the center of gravity of the half-sphere’s area. We thus find a volume

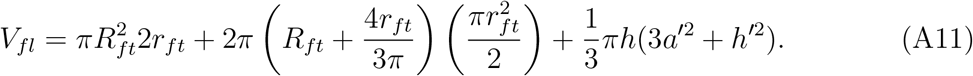

Note that the volume of the spherical caps can be negative if the cap is concave (*h <* 0).

The surface area is given by the surface area of the spherical caps (irrespective if concave or convex) plus the surface area of the “half-torus”, which is calculated again following Guldinus second theorem by rotating a half-circle around the radius between the axis of rotation and the center of gravity of the half-circle line. We thus find the surface area

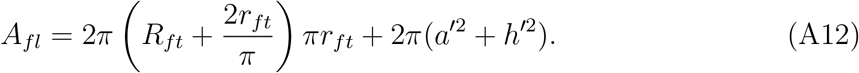

Due to the geometry, we have the additional constraint *h < r*_*ft*_ for concave spherical caps.

The mean squared radius within the torus formed by the filament bundle is

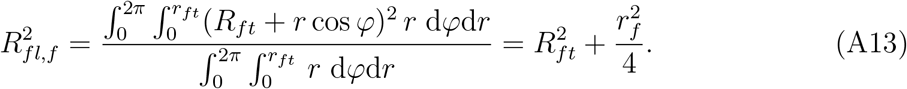

The torus with embedded filament bundle is modeled as a torus of the liquid phase engulfing the filament bundle as shown in Fig. A1 (d). The surface area and volume of the initial torus shape still apply at the condition that 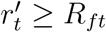. The mean squared radius within the torus is the same as for the filament-embedded lens shape.

#### b. Minimizing free energy

In order to minimize the energy of the droplet-filament system, we consider the surface energy *E*_*surf*_ of the droplet and bending energy *E*_*bend*_ of the filament. The surface energy is given as

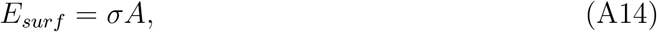

with surface tension *σ* and surface area *A*. The bending energy is

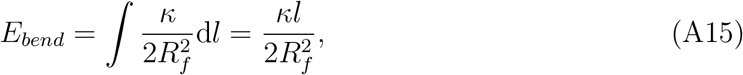

with bending rigidity *κ*. For the second model, using a bundle of filaments instead of a single overlapping filament, the bending energy needs to be adjusted to account for the packing fraction *ε*. For this model we follow [19] and use

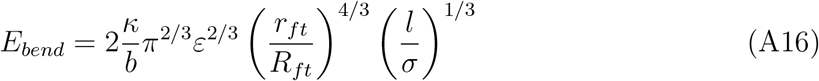

for the calculation of the bending energy. Here, *σ* is the diameter of the filament and *ε* is the volume fraction of the filament inside the filament bundle.

Since we cannot determine *ε* from experiment data, we turn to our simulations. From the cross sectional data from the simulations with the non-overlapping filaments (see. Supplement Fig. S5(e)) we can see that the distance between filaments inside the filament bundle is at around one particle. This gives, assuming no underlying packing structure, a packing fraction of *ε* = (*σ/*(*σ* + *σ*))^2^ = 0.25, where the denominator is the size of the filament including a wetting film. Assuming now slightly higher packing due to different effects such as not perfect sphericity of the filament bundle we end up with *ε* = 0.27.

The effective energy of the system is then the sum of the bending and surface energies. We thus have three input parameters: *σ, κ* and *l*. An additional parameter is hidden in *R*_*f*_ and is given by the radius of the initial sphere *R*_0_, which sets the base curvature and thus modifies the magnitude of *E*_*bend*_.

As seen above, both lens and torus geometry can be reduced to one varying parameter (*h* for the lens and *r*_*t*_ for the torus) that links *R*_0_ and *R*_*f*_ for the respective shape. Thus, we can calculate the energy landscape for each geometry as a function of *R*_*f*_ if *R*_0_ is known. Energy minimization with regard to the free parameter, however, requires solving higher order polynomials. Instead, we calculate plausible shapes and their energy and thereby find the minimum energy shape for each geometry, see Fig. 2 (a). Upon introducing additional constraints for the kettlebell and shapes containing a filament bundle, energy minimization is done numerically using SciPy [38].

Incorporating the kettlebell as a possible shape in our analytical model requires another parameter: the film thickness *r*_*film*_. For the first analytical model, *r*_*film*_ constraints the possible geometries of the kettlebell and the torus. For the torus it sets a maximum value for *r*_*t*_ while it fixes *r*_*t*_ = *r*_*film*_ in the case of the kettlebell. Based on the simulation results, we set *r*_*film*_ = 60 nm.

In the second model (shapes containing a filament bundle), we further need the volume fraction *ε* of filament within the filament torus. Leaving *r*_*film*_ and *ε* unconstrained during the minimization results in *r*_*film*_ → 0 and *ε* → 1. Indeed *r*_*film*_ becomes obsolete, as the kettlebell is always the filament torus with additional sphere and the additional film in the torus geometry is unconstrained. However, *ε* → 1 opposes the assumptions such that we set *ε* = 0.27, as mentioned above.

For filaments long enough to conform to *R*_*f*_, i.e. larger than 2*πR*_*f*_ for a given shape, we can calculate the total energy for each shape as a function of *R*_*f*_. Minimal energy morphologies with shorter filaments are likely not solids of rotation and not considered here. A minimal filament length is therefore required and some geometries become inaccessible.

Morphologically, the sphere shape can transform smoothly into a lens shape but has to change topology when transitioning into a torus. This transition can be subject to an energy barrier, see Fig. 2 (a), such that a system may settle in the first minimum given by the lens shape.

The central assumption is that the droplet is above the bendocapillary length, i.e. that the adhesion energy between droplet and filament is larger than the bending energy to hold the filament within the liquid phase.

### 3. Calculation of the bendocapillary length

The bendocapillary length *R*_*bc*_ defines the radius of a radius at which the wetting energy of a contacting or embedded filament and its bending energy needed to comply to the geometry of the droplet are equal. Consequently, droplets with a radius exceeding *R*_*bc*_ are expected to contain the bent filament, whereas droplets smaller than *R*_*bc*_ cannot retain the filament.

As above, the bending energy of the filament bent to a radius *R*_*max*_ is

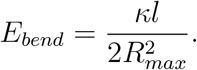

A typical measure for rigidity in polymers/filaments is the persistence length *l*_*p*_, which is the length over which the local orientation of the filament becomes uncorrelated [39]. *l*_*p*_ can be calculated as

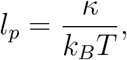

with the energy of thermal fluctuations *k*_*B*_*T*.

The adhesion energy *W* of a filament with the droplet phase is the difference in energy the filament experiences upon coming into contact with the droplet. Using Young’s equation we can relate the surface tension between filament and buffer *γ*_*bf*_, between filament and droplet *γ*_*df*_ and between buffer and droplet *γ* with contact angle *θ*

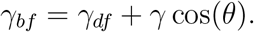

We assume a filament with a high affinity for the liquid phase as observed, e.g. for PLPPR3 and actin. We thus assume a contact angle of *θ* = 0° and find *W* (*θ* = 0°) = *A*_*contact*,0°_(*γ*_*bf*_ − *γ*_*df*_) = 2*πr*_*f*_ *lγ*, with filament radius *r*_*f*_. Note that at perfect wetting *γ*_*bf*_ ≥ *γ*_*df*_ + *γ*, which is equal only at the limit *θ* → 0°. Higher differences in surface tension between the phases are also possible.

Equating bending and wetting energy, we then find

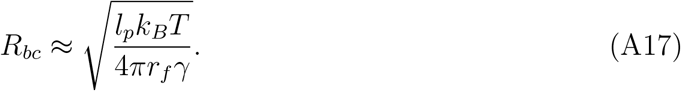

Note that neutral wetting conditions (but assuming the same affinity between filament and droplet phase, *θ* = 90°) lower *R*_*bc*_ by a factor of 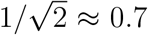 due to changing contact area, whereas yet higher affinity of the fiber to the droplet phase increases *R*_*bc*_.

**TABLE 1.**
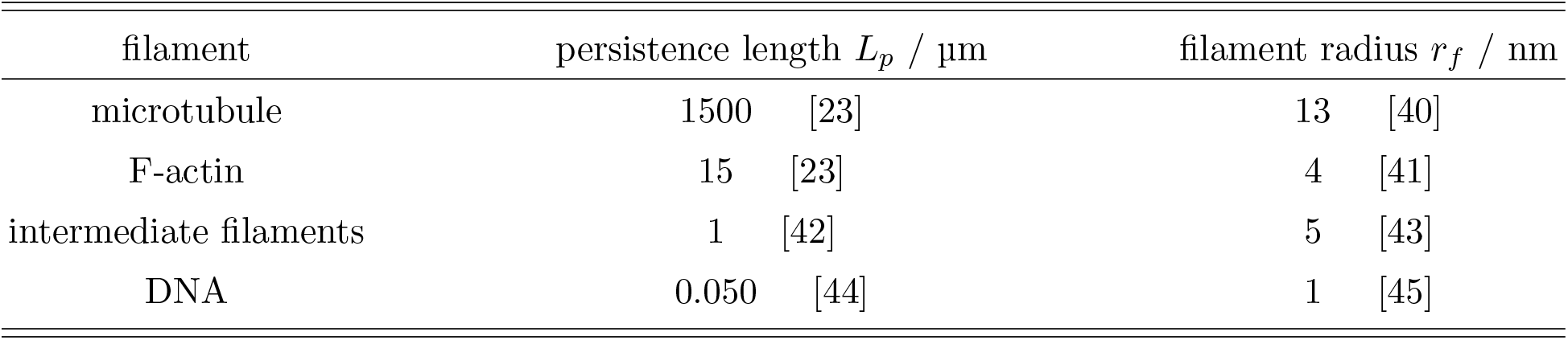
Material properties of different biopolymers used to estimate *R*_*bc*_.

### 4. Simulations

#### a. Simulation setup

To capture the nature of the experimental system, we built a simulation environment consisting of two types of particles, droplet and filament particles. The droplet particles phase-separate spontaneously and form the condensate and a surrounding gas phase in the simulation. The filament particles are assembled into one long fiber.

Droplet particles interact with each other via the truncated and shifted Lennard-Jones potential:

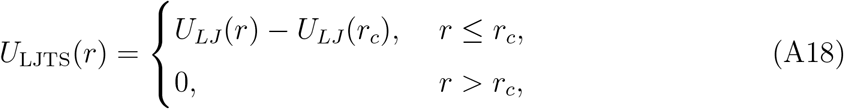

where

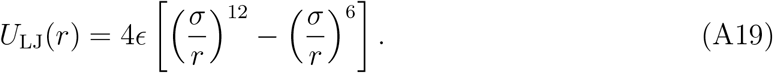

Between the droplet particles, the potential functions as an attractive force that gives the particles the ability to spontaneously phase-separate on their own above a critical concentration. Using a depth of *ϵ*_*ll*_ = 1*/*0.65 *k*_*B*_*T* for the potential well, a size of *σ* = 0.03 *µm* for the beads and a cut-off radius of *r*_*c*_ = 2.5*σ* the particles form a droplet with a surface tension *γ* = 0.448 *ϵ*_*ll*_*/σ*^2^ = 2.945 · 10^−6^ J/m^2^, calculated following Irving-Kirkwood [46]. The surface tension is in the range of previously reported values of biological condensates [22, 31].

To model the filaments inside the droplet we use only a single filament instead of having multiple filaments of varying length. We use this simplification as we are mostly interested in the interplay of surface tension of the droplet and bending rigidity of the filament and a single longer filament should approximate the sum of the bending energies of multiple smaller filaments. The filament is modeled as a chain of self-avoiding beads. The length extension of the filament is controlled by harmonic springs between the individual beads,

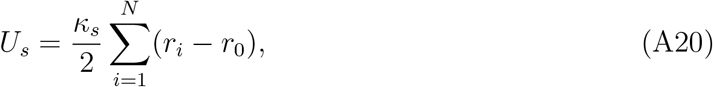

where *κ*_*s*_ = 0.17 *N/m* is the spring constant and *r*_*i*_ = *σ* is the equilibrium distance. The result is a filament with minimal length fluctuation during simulations, while still stretching more than actin filaments, with a strain of 0.012 under a force of 60 pN in the simulated fiber compared to 0.002 for actin [47]. However, higher values of *κ*_*s*_ would require smaller time steps Δ*t* during the simulation. We checked our simulations and confirmed that no significant extensions occur during our simulation that would warrant a higher spring constant and therefore a lower Δ*t*. To control the bending rigidity of the filament we use harmonic angular springs between the beads given by

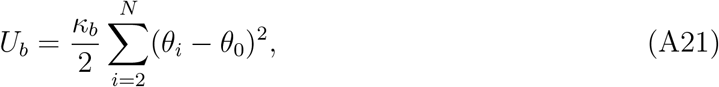

where *κ*_*b*_ = 3.34·10^−18^ Nm (corresponds to 1·10^−25^ Jm in the continuum model, actin should have *B*_*S*_ = *E*_*y*_*I* = 2.12 · 10^−25^ Jm, Young’s modulus *E*_*y*_ ~ 1.8 GPa = 1.8 · 10^9^ N/m^2^ [47] and actin radius 3.5 nm (*I* = *πr*^4^*/*4)) is the spring constant and *θ*_*i*_ = *π* is the equilibrium angle. This provides a filament which prefers to be straight and has a similar bending rigidity as F-actin.

Further, an attractive potential between filament and fluid particles is required for the filament to be incorporated into the droplet. This interactive potential represents the wettability of the filament and is modeled using the LJTS Eq.A18 with *ϵ*_*lf*_ = 1*/*0.6 *k*_*B*_*T, σ* = 0.03 µm and *r*_*c*_ = 2.5 *σ*. With the structure and wettability of the filament defined, we create two different models by tuning the interaction potential of non-sequential filament particles.

The simulations are performed in a 2 × 2 × 2 µm^3^ simulation box with periodic boundary conditions. It evolves according to Langevin dynamics in the NVT (canonical) ensemble at *T* = 300 K. For the Langevin dynamics, a damping factor *τ* = 2480, particle mass *m* = 700 pg and time step Δ*t* = 2 µs are used providing a diffusion constant *D* = *k*_*B*_*T/*(3*πηd*) at times much larger than *τ*, where *η* = 0.001 Pas [48, 49] is the viscosity of cytoplasm and *d* = 0.03 µm is the diameter of the particle [50].

To initialize the overlapping fiber simulations, the filament is constructed into a loop of radius *R*_*f*_ with initially non-overlapping particles, placed inside the simulation box, and made rigid. Next, droplet particles are distributed randomly inside the simulation box and allowed to condensate and equilibrate around the filament for 8 · 10^6^ time steps, resulting in a liquid phase engulfing the filament. Afterwards, the rigidity constraint of the filament is removed and the system is relaxed for another 2 · 10^7^ time steps where both filament and droplet particles can move freely. Note, however, that the filament is already kinetically trapped and barely moves.

To initialize the non-overlapping fiber simulation, the filament is placed in the simulation box in the form of a small rigid coil of diameter ~ 240 nm (8 *σ*) and the fluid particles are randomly distributed in the simulation box around the filament. For the first 8 · 10^6^ time steps the droplet particles are allowed to equilibrate and condensate around the filament. Next, the rigidity constraint of the filament is removed and the bending coefficient is stepwise increased (8 steps à 2 · 10^6^ time steps) until the final bending energy of *κ*_*B*_ = 3.34 · 10^−18^ Nm is reached. Finally, the system is allowed to relax for another 1.2 · 10^8^ time steps.

#### b. Particle packing

For spheres the densest packing is achieved by a hexagonal lattice. The number of particles surrounding a single particle in the middle of an hexagonal lattice is given by

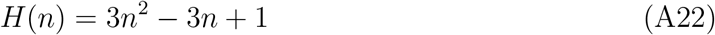

where *n* is the *n*th hexagonal number. Assuming that our filament is the central sphere in this case, we can calculate the expected hexagonal number. For our truncated and shifted Lennard-Jones potential with *r*_*C*_ = 2.5*σ* the expected hexagonal number would be *n* = 2. Indeed, if we plot the expected number of particles from hexagonal packing as a function of filament radius, we can see that fit aligns completely with the border region between the ring and the kettlebell, see Supplementary Fig. S4.

#### c. Data analysis simulations

Data analysis of the simulation data is performed using Python with the packages NumPy [51], SciPy [38], pandas [38, 52] and scikit-learn [53].

To identify droplets in the simulation we used scikit-learn’s DBSCAN. The clustering was performed on a euclidean distance matrix with *ϵ* = 2*σ* and min samples=20.

To determine the surface tension of the droplet in the coarse-grained molecular dynamics simulations, we followed the method presented by Thompson et al. [54] based on the theory of Irving-Kirkwood [46]. The surface tension can be entirely determined as function of the radial distance from the center of the droplet *r*. We used a system with 10^4^ particles and let it equilibrate for 2 · 10^7^ time steps. The normal component of the pressure tensor is given

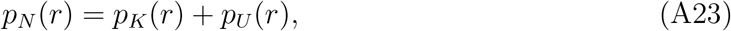

where *p*_*K*_(*r*) = *k*_*B*_*Tρ*(*r*) is the kinetic term and *p*_*U*_ (*r*) = *S*^−1^ ∑_*k*_ *f*_*k*_ is the configurational term. *k*_*B*_ is the Boltzmann constant, *T* is the temperature, *ρ*(*r*) is the density function, *S* = 4*πr*^2^ is the surface area of a spherical surface of radius *r*, and *f*_*k*_ is the normal component of the interparticle components across the surface *S*. We can then determine the surface tension using

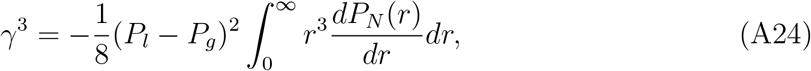

where *P*_*l*_ and *P*_*g*_ are the liquid and gas pressure, respectively, far from the interface.

To create the shape phase space for the simulations (Fig. 3(b) and 2(b)), a support vector machine from scikit-learn [53] was trained on the respective simulation data. To achieve the best possible result without bias we used optimized the parameters for the support vector machine following [55]. The trained model was then used to classify tuples of droplet volume and filament length as spheroid, kettlebell or torus. The results of this classification are presented shown in the figure.

## Appendix B

**Supplementary Figures**

**FIG. S1.**
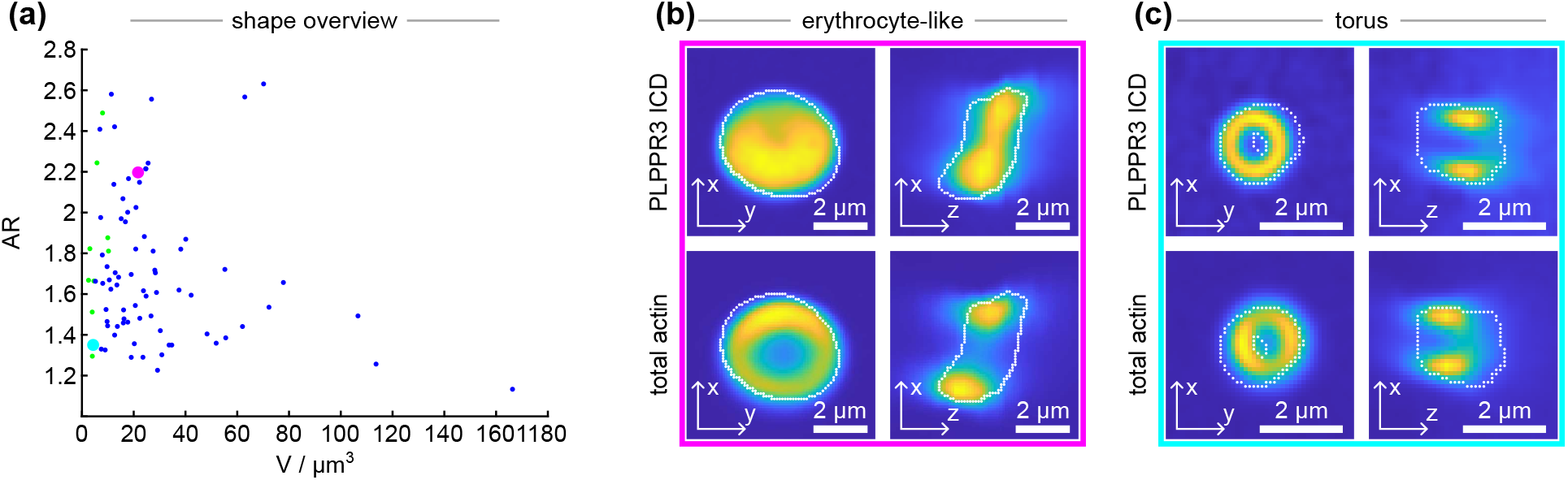
Analysis and confocal images (dual-camera mode) of PLPPR3 ICD condensates with F-actin observed with 4 µM actin and 10 µM PLPPC3 ICD. (a) Aspect ratio as a function of droplet volume from N=82 condensates. Blue circles indicate spheroid, green circles indicate ring morphologies. (b)-(c) The same condensate as in Fig. 1(e)-(f) but with detected outline indicated by white dots. The condensate is highlighted in magenta and cyan respectively in panel (a). Note that the channels in the dual-camera mode are not fully aligned leading to an inflation of recognized condensates and resulting lower AR.

**FIG. S2.**
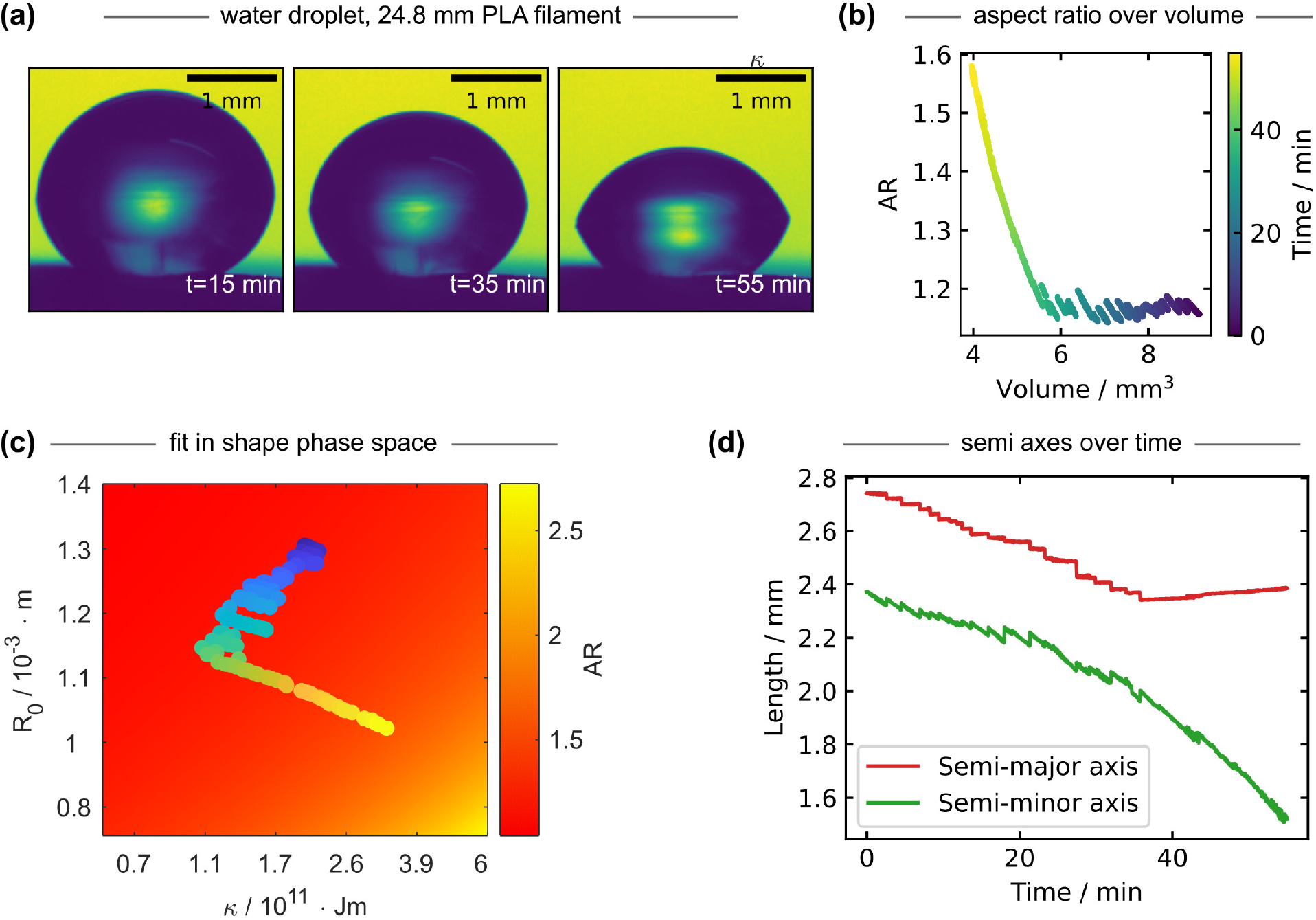
Experiments with water droplets engulfing a PLA filament match the analytical model. (a) Images of the droplet with embedded filament on a hydrophobic taro leave taken over time as the droplet evaporated. (b) Time evolution of the semi-major and semi-minor axis of the water droplet containing a PLA filament depicted in Fig. 1. (c) Fitting experimental data to the shape-diagram created from the analytical model for the water droplet containing PLA filament experiment can be used to determine the bending rigidity *κ*. Filament length *L* = 24.788 mm and surface tension *γ* = 0.072 J/m^2^ are fixed to known values. Droplet volume and *κ* are free parameters. Volume and AR are aligned with values from analytical model. Color gradient shows time as in panel (b). Based on supplier data, the filament has a Young’s Modulus of *E*_*y*_ ≈ 3.54 GPa and optical measurements show a filament radius of 12 to 13.5 µm. We thus expect *κ* between 5 and 9 · 10^−11^ Nm^2^. Variations in *κ* throughout the shape relaxations may be caused by effects of gravity at early times and friction in the filament at late times. Moreover, we expect contributions of wetting to the substrate throughout the experiment. (d) the time course of the major and minor axes of the shape reveals discrete steps in the major axis, suggesting static friction between the filament loops.

**FIG. S3.**
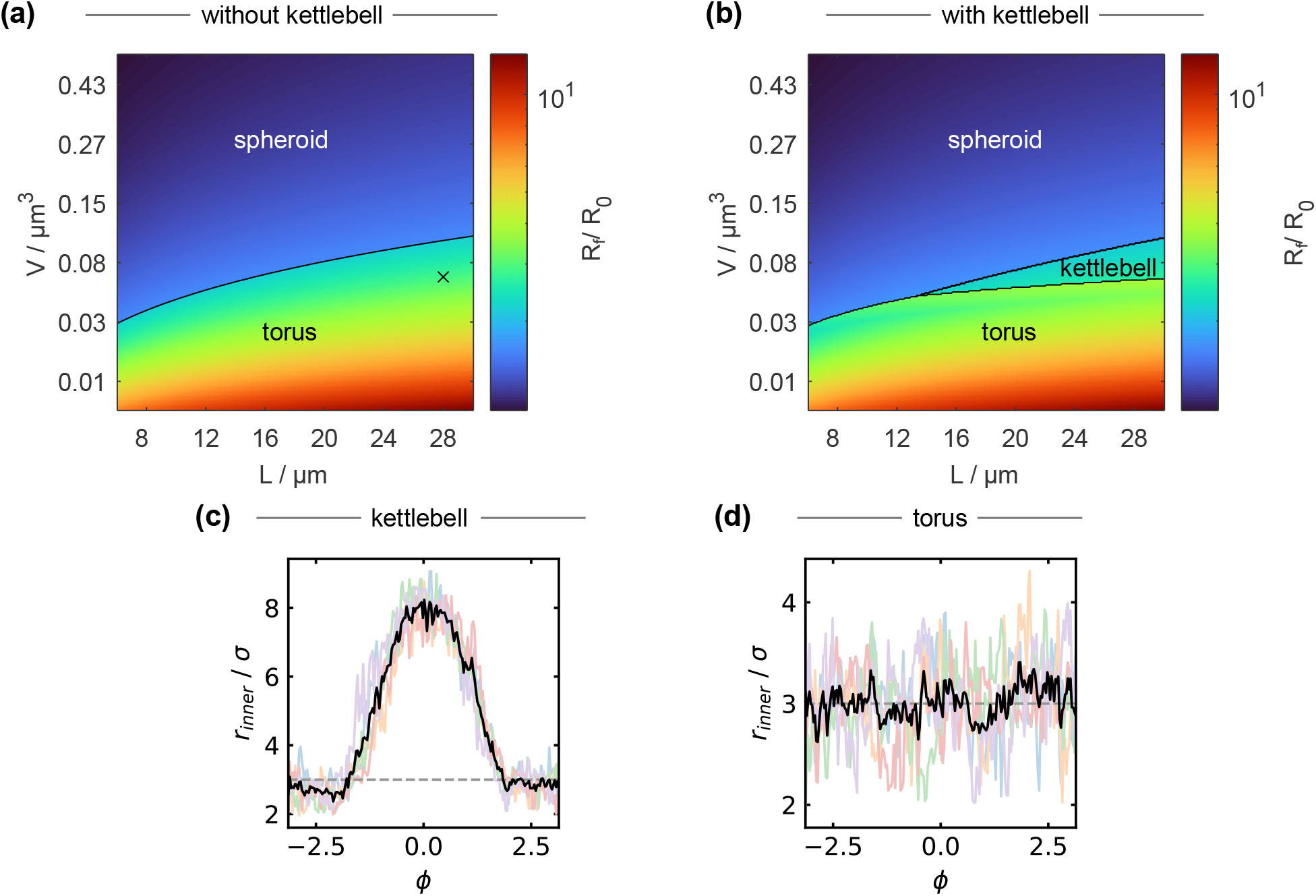
Introducing the kettlebell shape, i.e. a constraint on the wetting film of the torus, changes the shape space. (a) Shape space for varying *V* and *L* with spheroid and torus as possible shapes with surface tension *γ* = 1 · 10^−5^ J/m^2^ and bending stiffness *κ* = 1· 10^−25^ Jm. (b) Shape space with the same parameters as (a) but with kettlebell as additional possible shape. The film thickness is set to *r*_*film*_ = 60 nm to be analog to the our observations form the simulations. (c),(d) radial distributions of the minor radius of a kettlebell and torus, respectively along the azimuthal angle *ϕ*. This distribution is used to discern between tori and kettelbells.

**FIG. S4.**
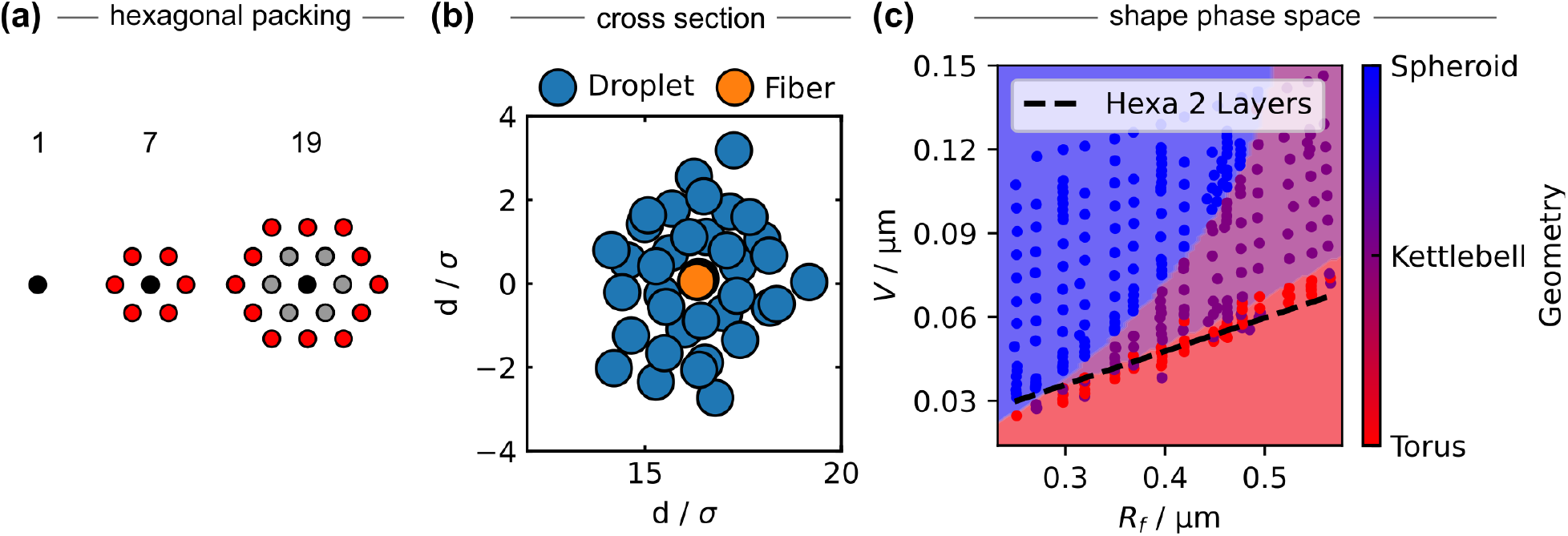
Packing of droplet particles around the filament in torus configuration. (a) Theoretical packing of spheres in a hexagonal lattice. (b) Cross section of a torus highlighting the structured droplet particles around the filament. (c) Shape diagram with line that shows the theoretical volume of a torus of a radius assuming hexagonal packing of droplet particles around filament particles.

**FIG. S5.**
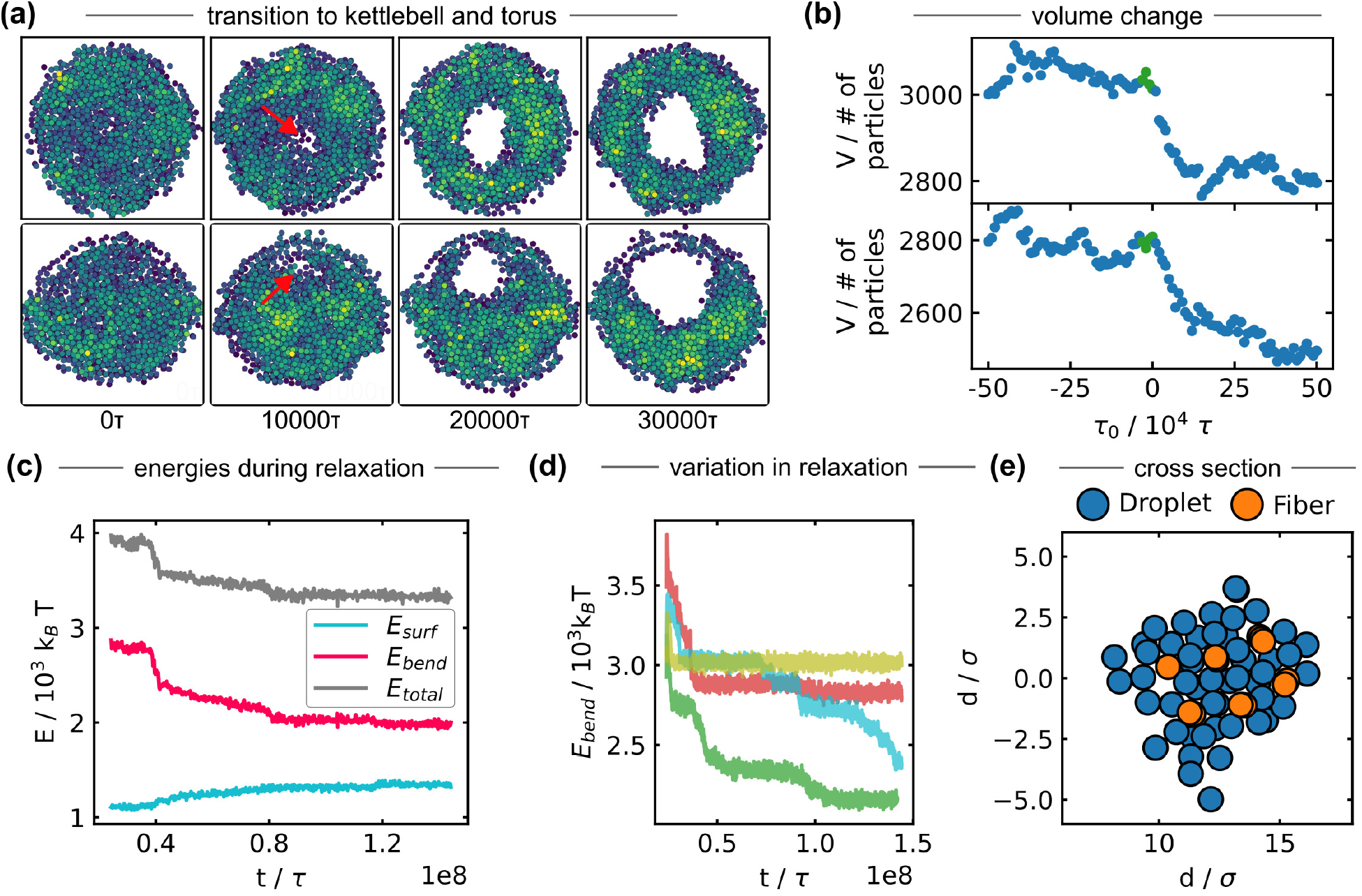
The transition from spheroid to torus or kettlebell happens at a short time scale. (a) Top-down view of the transitioning process from a spheroid to a torus (top) and kettlebell (bottom). (b) Volume of droplet before and after the transition. The green dots indicate the position of the images on the left. *τ*_0_ is the point at which the transition is fist observable. (c) Trajectories of the bending energy *E*_*bend*_, the surface energy *E*_*surf*_ and the total energy *E*_*total*_ during the relaxation periode for an example simulation (*n*_*part,liquid*_ = 4500 and *n*_*part,filament*_ = 400). (d) Four different trajectories of the relaxation of the bending energy for the same simulation system but with different random seeds (*n*_*part,liquid*_ = 4000 and *n*_*part,filament*_ = 350). (e) Cross section of a torus showing the ordering of the filament inside.

**FIG. S6.**
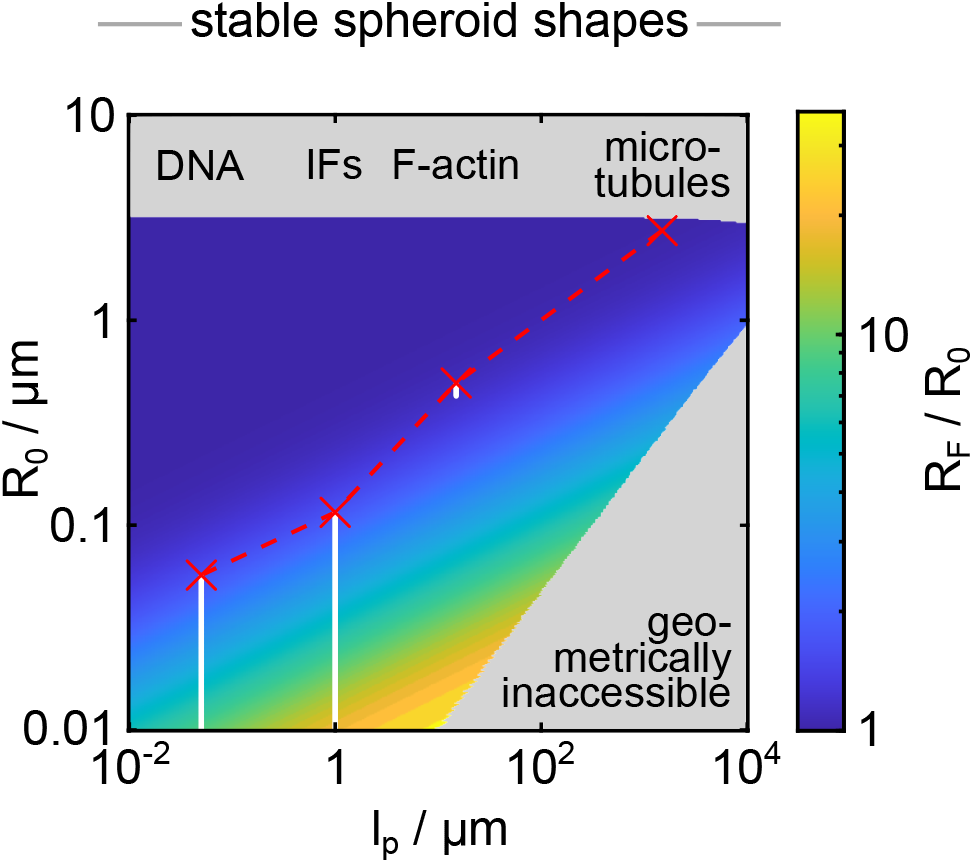
Mutual deformations of biomolecular condensates and different biopolymers expand the regime in which droplets retain the filament(s), also considering spheroid shapes only. Shape diagram of minimal energy spheroid shapes for a condensate with a surface tension of 5 · 10^−6^ J/m^2^ and filament length of 20 µm as a function of condensate volume and filament bending rigidity. The bending rigidity is expressed in terms of persistence length and the filament volume in terms of radius of a sphere of the same volume *R*_0_ for convenience. The colormap shows the aspect ratio of the minimal energy shapes as *R*_*f*_ */R*_0_. The red crosses indicate the approximated *R*_*bc*_ for microtubules, F-actin, intermediate filaments and DNA. All shapes above this line are stable. The white dots indicate additional shapes stabilized by mutual deformations. See Fig. 5 (a) for full shape space considering also torus shapes.

